# Interpretable per Case Weighted Ensemble Method for Cancer Associations

**DOI:** 10.1101/008185

**Authors:** Adrin Jalali, Nico Pfeifer

## Abstract

**Motivation:** Molecular measurements from cancer patients such as gene expression and DNA methylation are usually very noisy. Furthermore, cancer types can be very heterogeneous. Therefore, one of the main assumptions for machine learning, that the underlying unknown distribution is the same for all samples, might not be completely fullfilled. We introduce a method, that can estimate this bias on a per-feature level and incorporate calculated feature confidences into a weighted combination of classifiers with disjoint feature sets.

**Results:** The new method achieves state-of-the-art performance on many different cancer data sets with measured DNA methylation or gene expression. Moreover, we show how to visualize the learned classifiers to find interesting associations with the target label. Applied to a leukemia data set we find several ribosomal proteins associated with leukemia's risk group that might be interesting targets for follow-up studies and support the hypothesis that the ribosomes are a new frontier in gene regulation. **Availability:** The method is available under GPLv3+ License at https://github.com/adrinjalali/Network-Classifier.

## 1 Introduction

Over the past decades, biology has transformed into a high throughput research field both in terms of the number of different measurement techniques as well as the amount of variables measured by each technique (e.g., from Sanger sequencing to deep sequencing) and is more and more targeted to individual cells ([23]). This has led to an unprecedented growth of biological information. Consequently, techniques that can help researchers find the important insights of the data are becoming more and more important. Predicting survival of cancer patients based on measurements from microarray experiments was first proposed more than a decade ago, but there is often very little overlap between the important genes or biomarkers identified by different studies ([8]). Several reasons have been suggested that could explain these findings (e.g., heterogeneity of cancer samples or insufficient sample size). Attempts have been made to incorporate additional information from other sources such as protein-protein interaction (PPI) networks to make the predictions more robust ([5]). One of the latest approaches integrated network and expression data by introducing a network-induced classification kernel (NICK) ([17]). Although this method was shown to have state-of-the-art performance, it is not straightforward to determine which are the important variables that led to the good performance. Furthermore, a central assumption underlying many of the methods is that all data are drawn from the same unknown underlying distribution. Especially for heterogeneous cancer samples this might not be the case, in particular not for all measured genes.

In this work, we introduce a method that is aware of this potential bias and utilizes an estimate of the differences during the generation of the final prediction method. For this, we introduce a set of sparse classifiers based on *L*1-SVMs ([3]), where each set of features used by one classifier is disjoint from the selected feature set of any other classifier. Furthermore, for each feature chosen by one of the classifiers, we introduce a regression model based on Gaussian process regression that uses additional features. These regression models are then used to estimate how predictable features of each classifier are, for each test sample. This information can then be used to find a confidence weighting of the classifiers for the test sample (up-weighting classifiers with high confidence and down-weighting classifiers with lower confidence). Schapire and Singer showed that incorporating confidences of classifiers can improve the performance of an ensemble method ([22]). However, in their setting confidences of classifiers are estimated using the training data and are thus fixed for all test samples, whereas in our setting we estimate confidences of individual classifiers per given test sample. We show that this method has state-of-the-art performance on several gene expression data sets as well as for methylation data sets both for different cancer types. Since the weighting of the classifiers is customized for each test sample the estimated confidences can offer insights into the specific characteristics of the cancer of each individual. Therefore, we show how to visualize the important genes found through such an analysis and then also show how to find the important genes of the training set using our learning method.

## 2 Methods

### 2.1 Materials

#### 2.1.1 Data Sources

In this article, our method is applied to two different data types: gene expression data and DNA methylation data.

- *Van ’t Veer* ([27]): This data set contains 97 breast cancer samples. For each sample gene expression measurements as well as prognosis labels are available, indicating if the patient had any distant metastases after diagnosis.
- *The Cancer Genome Atlas* ([24]): The Cancer Genome Atlas (TCGA) is a joint effort of the National Cancer Institute and the National Human Genome Research Institute to advance the understanding of the molecular basis of cancer. They provide access to the different measurements from cancer samples that have been analyzed to external researchers. Samples are categorized according to diagnosed cancer from which we use the following groups:

- *Acute Myeloid Leukemia (LAML)* ([26]): At the time of writing, the data set includes 200 samples. 194 samples contain methylation data and we use the part of the data measured by JHU-USC HumanMethylation450 arrays. 173 samples contain mRNA data measured by HG-U133 arrays. In this article the methylation data is referred to as TCGA-LAML. Among available characteristics of samples, “risk group” and “vital status” are chosen as target classes. These labels show the aggressiveness of the disease. In our analysis, regarding risk group, {favorable} and {intermediate/normal, poor} samples form our two group, and in the analysis of vital status, {alive} and {dead} samples form our two groups of samples.
- *Breast invasive carcinoma (BRCA)* ([25]): This data set includes 993 samples with clinical data, and we use the methylation data component measured by JHU-USC HumanMethylation450 arrays. Only very few samples in this data set are indicated as having metastasized (8 samples). Therefore, it is not possible to analyze the data set according to metastasis as is done with van ’t Veer data. Hence the data are analyzed according to “tumor size”, “affected nearby lymph nodes”, “stage”, and “estrogen receptor”. Estrogen receptor was shown to be an important factor in prognosis ([16]), and along with other factors directly affects the decision for therapy ([12, 19]). For tumor size {T1, T2} samples are one category and {T3, T4} the other category; in order to analyze affected nearby lymph nodes, {N0} is compared to {N1, N2, N3}; stage is analyzed as having {stage I, stage II} vs. {stage III} samples. Estrogen receptor status of samples is either positive or negative, and they form our two classes.

#### 2.1.2 Data Preprocessing

To prepare gene expression data for analysis, microarray probes are mapped to their respective gene. If there are multiple probes for a gene, the median reported gene expression value of those probes is adopted as the gene expression for that gene.

Preparing the methylation data, we use the nearby gene for each methylation site available for each sample and each methylation site. The median beta value of methylation sites mapped to each gene is taken as the methylation value of the corresponding gene. In this process only methylation sites located on the promoter region of a gene are considered and others are discarded.

### 2.2 Boosting

For a given prediction problem the idea of boosting is to find an optimal combination of classifiers, also called “weak learners” ([6]). There are many methods of finding the optimal combination of such weak learners, two of which are stochastic gradient boosting ([10]) and AdaBoost ([9]). Stochastic gradient boosting tries to estimate the gradients of the loss function and train each individual weak learner in a way that best improves the loss function. AdaBoost tries to identify samples among given data samples that are harder to classify, and gives them more weight in the process of training individual weak learners. One way of improving AdaBoost is to take into account the confidences of predictions given by weak learners if possible and use estimated confidences in the voting process ([22]).

### 2.3 Learning a Mixture of Disjoint Classifiers

When dealing with cancer, we need to consider the fact that tumors of the same type of cancer can be very different in nature and they are usually classified as different cancer subtypes. In fact, even one single tumor can be very heterogeneous ([13]). This means that the malignancies causing the cancer to happen are genetically different between subtypes, or even within subtypes, and it is possible to have multiple underlying cellular processes causing a particular cancer. Also it is important to note that the nature of our given data is such that the input features are properties measured from genes, e.g. gene expression or methylation values, and these variables are correlated and statistically dependent on each other. Our method tries to exploit these properties of the problem to infer an interpretable model with state-of-the-art performance.

Our method can be characterized by the following key parts:

Training phase:

- Fit several individual classifiers to the data, in such a way that the features of the data they use are disjoint sets.

Prediction phase:

- Calculate the prediction confidence of each individual classifier by:

- Estimating the reliability of input features of the classifier;
- Estimating the confidence of the output based on the decision values.
- Calculate a weighted prediction label based on the individual classifier confidences.

#### 2.3.1 Properties of the Individual Classifiers

A wide variety of classifiers is possible within our framework. One requirement is that the classifier is regularized (i.e., the stronger the regularization, the less complex the model gets and consequently the less features are used). The classifier is also required to report the probability of its calculated output, or to give a decision value according to which it chooses the predicted class. We use an *L*1 regularized SVM for this purpose with a linear kernel ([3]). The *L*1 regularization makes the SVM sparse, i.e. using only a few input features, and the linear kernel allows us to infer which features are used in the decision function of the SVM after it is fit to the data.

#### 2.3.2 Training the Individual Classifiers

The model starts with no individual classifiers and an empty set of excluded features. In each step, the excluded set of features is removed from the data, then a classifier is fit to the data. Next the features used by the most recent trained classifier are added to the excluded set. In the case of a linear kernel SVM, this is achieved by finding features with a non-zero coefficient in the model. This way the features being used by classifiers are disjoint and might represent different underlying causes of groups into which samples are to be classified.

#### 2.3.3 Combining Classifiers by Estimating Confidences of Individual Predictors

Given a set of classifiers, the question is how to combine them to come up with a joint prediction value for each test sample for which we want to predict the output label. The intuition behind combining the classifiers is to put more weight on classifiers that use features whose behavior is similar to the training data. This is motivated by the fact that some parts of the test data might behave very differently to the training data, meaning that a classifier using these features should have lower performance than a classifier using features that are distributed similarly to the training data. Therefore we need to evaluate the reliability of the input features of each individual classifier. In scenarios like gene expression or methylation analysis, we usually have many input features. Furthermore, many features are correlated and statistically dependent. The idea of our new method is to build separate prediction models for each feature of each classifier. These prediction models can then be used to obtain a confidence for the feature in a given test sample. These confidences can then be combined for each classifier to give a weighting of the classifiers for the given test sample. To evaluate an observed feature *f*, we try to choose a few statistically dependent features, and fit a model to predict *f*. To find these features, first the estimated mutual information of all other features with feature *f* is calculated ([21]). Then, features having mutual information value within the top 5%, or at least 5 features, are selected as predictors of *f*. Given a test sample, the closer the predicted value of *f* is to the observed value, the more reliable it is. To quantify this, we need to not only know the predicted value of the feature, but also a confidence interval for that prediction. This can be achieved using Gaussian processes, which give the mean and variance of the posterior probability under the condition of observed values for selected features. A weighted average of these values gives us the overall reliability of the features of an individual classifier.

In addition to the confidence in the classifier estimated by looking at the confidences of its individual features, we also account for the confidence that the classifier has in the prediction label of the test sample. If the method supplies such a confidence value (e.g., Gaussian processes), we can directly use it. Otherwise, we estimate it using the decision value. In our setting, the linear SVM gives a decision value whose sign defines the predicted class. Using these values we estimate a confidence for each individual classifier. Several approaches exist for deriving a confidence from the decision values ([18]). Whether these or other additional methods could lead to further improvements of our method, will be topic of further study.

More formally speaking, define *X* to be the set of input samples, *X_s_* to be the input vector of sample *s*, *y_s_* and *ŷ_s_* to be respectively the original label and predicted output of sample *s*, Δ to be the set of individual classifiers, *l_i_* to be an individual classifier, 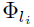 the set of input features of classifier *l_i_*, *l_i_*(*X_s_*) to be the label predicted by classifier *l_i_* for sample *X_s_*, and *f* to be a feature, *X_s,f_* to be the observed value of feature *f* in sample *X_s_*, 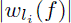 to be the absolute value of the weight of feature *f* in the decision function of classifier *l_i_*, and *g_f_* to be the Gaussian process predicting feature *f* using feature set Φ*_f_*. Also 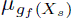 and 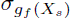 are the mean and standard deviation of the posterior probability given by Gaussian process *g_f_* under the condition of observing values of features in Φ*_f_*, and 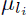 and 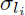 are respectively the expected mean and standard deviation of the decision value of classifier *l_i_*. Here *F* is the cumulative distribution function of a standard normal distribution.

The training phase of the model is shown in Fig. 1, in which, *N* is the number of individual learners to be included in the model, Φ*_l_* is the union over all 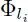 and 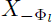 is the input *X* after discarding all features of the set Φ*_l_*. TOP is the function which selects the maximum of the top 5 and top 5% features *f*′ of all features ordered by mutual information with feature *f*.

**Figure 1:**
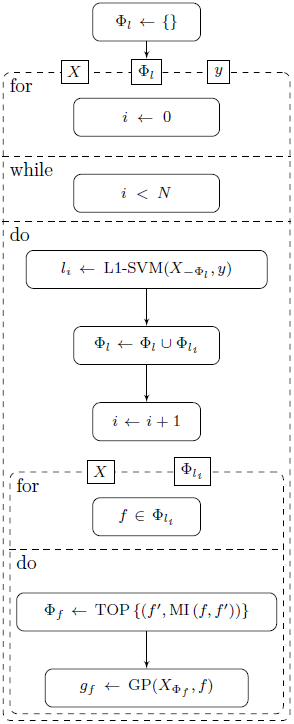
UML activity diagram of the training process

Now given a test sample *X_s_*, the estimated confidence of a feature *f* is:

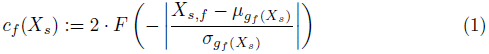

Then the overall feature reliability or confidence of a classifier *l_i_* is estimated as:

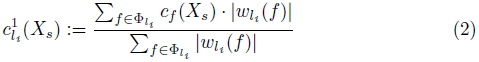

Also the estimated output confidence of the classifier *l_i_* is:

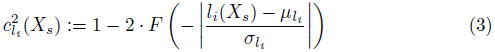

and the final confidence of the classifier *l_i_* is then:

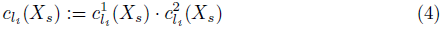

Finally, the predicted class *ŷ_s_* is calculated as the sign of a weighted vote among individual classifiers:

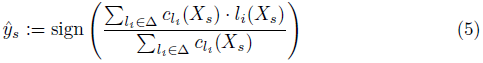

### 2.4 Visualization of Model Predictions

The interpretation of the model can be understood on two different ways. First we assume for a given training data set, the model is trained and a new test sample is given. For the given test sample it is possible to visualize the reliability of each used feature in individual classifiers, as well as the overall confidence of each individual classifier. Used features can be superimposed onto a PPI network as well as their reliability and the confidence of their respective individual classifier.

Gene expression and methylation level measurements from cancer samples are usually very noisy. Furthermore, cancers are usually very heterogeneous. Additionally, there might be different subgroups for each interesting group (e.g., cancer stage), for which the importance of the features also differs. To get a global picture of the important features, we therefore evaluate how often certain features are selected by the classifiers using 100 random train test partitionings with 80% of the data for training and 20% of the data for testing. To visualize high confidence relationships between features, we create a graph which has a node for every chosen feature in any of the 100 train partitions in any of the individual classifiers. The weight of an edge (*s, t*) is defined as the number of times the respective features have occurred together in an individual classifier. Then, all edges with low weights are discarded. In order to find a threshold to prune edges according to their weights, a Gaussian kernel density estimate is fit to the weights of the edges, and the threshold is chosen at the 90th percentile. Nodes that have an appearance frequency higher than the threshold are labeled by their gene names and edges having a higher weight than the threshold are kept in the graph.

For illustration purposes, choosing the regularization parameter is done in a way to maximize the number of genes selected with high confidence, as well as minimizing the number of genes pruned out in the process. It is important to remember that considering the results of the method under different regularization parameters is essential to make sure the selected genes possess a high confidence and are also stable regardless of sampling of the training data set.

### 2.5 Implementation Details

To compare the performance of our method with other methods, the implementations present in Python *scikit-learn(0.14)* package are taken. In the case of stochastic gradient boosting, the representing class is *GradientBoostingClassifier*, the number of classifiers is set to 100, and to make it sparse and prevent over-fitting, the maximum number of features for splits in trees is set to 5, and the maximum number of layers is set to 2. For AdaBoost, *AdaBoostClassifier* is used, which is an implementation of AdaBoost-SAMME ([30]), with weak learner set to *DecisionTreeClassifier* with maximum depth set to 2, and the number of weak classifiers set to 100. Parameters of the two boosting algorithms are chosen by a grid search on their parameter space over all the data sets and selecting the parameter sets which give a robust and stable result over all experiments. Detailed report of the performance measures with different parameters is given in Supplementary 2. As an SVM, *ν*-SVM with *ν* = 0.25 is used, once with a linear kernel, and once with an RBF kernel; *γ* parameter of the RBF kernel is set to (num of features)^−1^. The *ν* parameter is set to the maximum value for which the optimization function is solvable with *libsvm* for all analyzed data sets ([4]). Smaller values cause the SVM to overfit to the data and not generalize well. The Gaussian process's correlation function is a squared-exponential, and mutual information is estimated using *minepy* package ([2]).

The PPI network used in our analysis is from the Human Protein Reference Database (HPRD) ([20]). Almost all edges and relationships between proteins that are added to this database are manually extracted from literature by biologists, hence it has a lower rate of edges included in the database for which there is no evidence in the literature.

## 3 Results and Discussion

### 3.1 Performance comparison

The performance of the method was compared with that of two ensemble methods, AdaBoost and stochastic gradient boosting, as well as an SVM with linear kernel, and an SVM with an RBF kernel. We also included our implementation of the NICK method ([17]). We randomly partitioned the data into training and test sets with 80% of the data for training and 20% of the data for testing. To compare the performance of the different methods, Area Under the receiver operating characteristic Curve (AUC) ([7]) was calculated on the test set over the decision values returned by the methods on the individual samples. The process was repeated 100 times to reduce random effects. As seen in Fig. 2, overall performances of all methods are comparable. In some cases a single SVM works better, in some other cases ensemble algorithms give a better performance. However, in most cases an improvement in performance is observed by adding individual learners to the model, with the greatest gains due to the first few individual learners added to the model.

**Figure 2:**
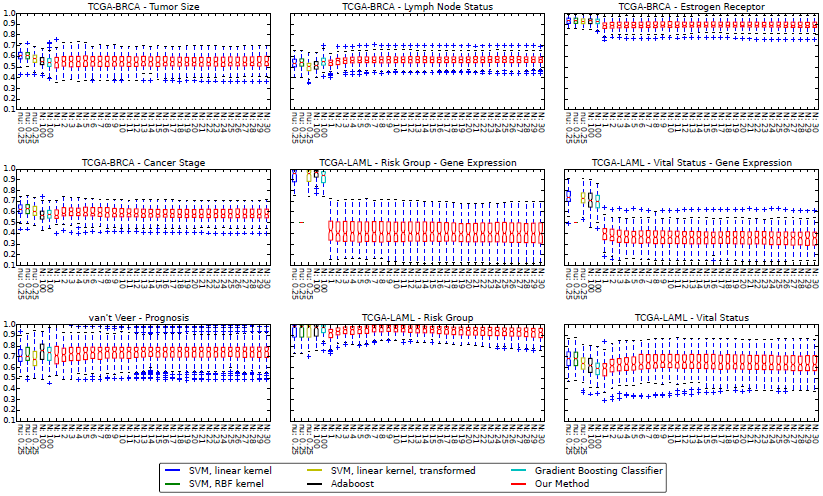
Performance Summary (AUC) Each box shows a 25–75% interval, as well as the median, which is shown as a horizontal line in each box.

### 3.2 Interpretability of Predictions

Here we present the results of running the method on the TCGA-LAML gene expression data set.

#### 3.2.1 Visualization of Features Important for a Particular Test Sample

Having a model trained on the data, and given a test sample, it is possible to infer and visualize which individual classifier(s) is (are) influencing the prediction most. To this end, individual learners as well as the features they use are visualized as in Fig. 3(a). In this figure, nodes with labels starting with “*L*___” represent individual classifiers, and other nodes are labeled with their respective gene name. The color of the node shows its confidence compared to other nodes; the darker the node, the higher the confidence. In the case of a gene, it is the confidence or reliability of the feature (*c_f_*), and in the case of an individual classifier, it is the overall estimated confidence (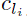). Edges show which classifier is using which genes in its decision function. The shape of a node represents the individual classifier they belong to.

**Figure 3:**
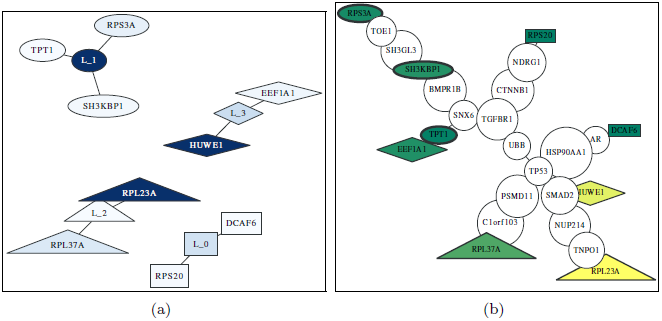
**Visualization of one model** A sample model for TCGA-LAML gene expression data **(a)** individual classifiers and their selected features; higher confidence of a node is shown by a darker color, **(b)** selected genes plotted over the PPI network; green and yellow show low and high confidence respectively, and the thickness of the border of the node shows the respective confidence of the individual classifier to which it belongs.

To get a better overview of the individual features that were chosen by the classifiers for the particular test sample, we visualized the corresponding genes on a graph containing information about the PPI network in Fig. 3(b). We extracted the PPI information from HPRD as explained before. This way, it is possible to find over-Rich No image in folderor under-regulated pathways that might be responsible for the label (e.g., cancer stage) of the test sample. Since PPI networks can be quite dense, we removed parts of the induced network. For this purpose we computed each shortest path between all pairs of selected features. Then, the minimum spanning tree of that section was plotted, after removing branches with no selected feature.

Most of the features chosen by any of the classifiers (colored nodes) are not connected to any other chosen feature. It is known that there is in many cases a correlation between expression value of the genes whose corresponding proteins interact ([14]). Therefore, a regularized model will only choose a subset of the correlated features. This explains the observation that features selected by a single model can be distant from each other on a PPI network; but if multiple disjoint sparse models are fit to the data, their selected features might happen to be close to each other on the PPI network (e.g., node TPT1 and node EEF1A1 in Fig. 3(b)).

#### 3.2.2 Visualization of Important Global Features

As explained in Section 2.4, a graph is created from model structures of all 100 random training partitions, and then it is pruned to keep only high confidence nodes and edges. The density estimation of the graph edge weights and the pruned graph are plotted in Fig. 4 and the nodes with labels are the ones that are not pruned. The nodes in this figure that do not have any label, are the ones with frequency lower than the corresponding threshold. Among the features considered to be important were features that had previously been linked to leukemia such as SH3KBP1 ([1]).

**Figure 4:**
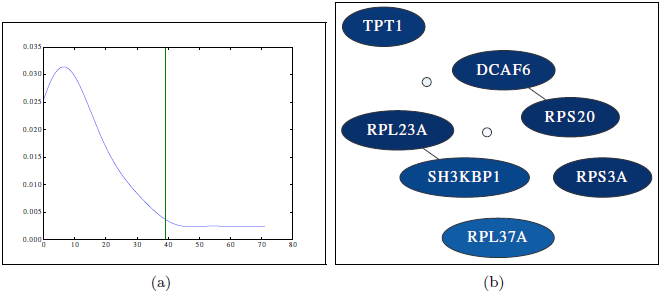
**(a) Determine pruning threshold** Threshold is determined by finding the point after which, 90% of the area under the curve is observed from left to right. The horizontal axis shows the observed frequency or weight of the edges. **(b) Important Global Features** High confidence nodes and edges of the graph generated from the model on TCGA-LAML gene expression data. Darker color represents higher rate of being selected by a classifier.

What was more intriguing to see was that four out of the seven important features of the TCGA-LAML gene expression data set contained ribosomal proteins (when using the risk group label). For a long time ribosomes were just considered machines that perform an unbiased translation of genes from mRNA to amino acid sequences, but this view has recently been challenged ([29]). One new hypothesis is that the ribosome introduces an additional regulatory layer. Therefore, it could very well be that mutations in ribosomal proteins can lead to a misregulation of expression levels of important genes and ultimately to the development of cancer (in this case leukemia). One of the ribosomal proteins we found was RPL23A. It has been shown that loss of RPL23A can impede growth and lead to morphological abnormalities in Arabidopsis Thaliana ([29]). Therefore, a mutation in RPL23A might also have severe effects in humans. A missense mutation in RPL23A was recently found in patients having Diamond-Blackfan anemia, which is an inherited form of pure red cell aplasia (related to leukemia) ([11]). Note that the model for LAML has low performance for the regularization value chosen. Nevertheless, the features shown here are also the ones with the highest confidence for models learnt with less regularization (with several other additional features). The models with less regularization show similar performance to the other methods shown in Fig. 2 (See Supplementary 1 for details).

## 4 Conclusion

Machine learning has become more and more popular in many real world scenarios for making sense of large collections of facts. Differences between the data used for training the method and new data for which the label should be predicted can limit the performance of prediction methods on those data. In this work we introduced a method that estimates these potential partial biases and incorporates them into the prediction function. We applied it to gene expression and DNA methylation measurements from cancer patients. Our method has state-of-the-art performance on many different prediction tasks. Further-Rich No image in foldermore, we show how to make sense of the predictions. Visualizing the important genes can lead to new biological insights, as shown for the TCGA-LAML data set with the risk group label. Instead of mapping the genes to PPI networks, one could also think of mapping them to signaling pathways ([15]). Recently, a study showed that most published signatures are not significantly more associated with cancer outcome than random signatures ([28]). One of the reasons for this finding is that the data comes from slightly different underlying hidden data distributions. Since our new method estimates this bias and corrects for it by up-weighting the classifiers that have higher confidence, we expect that it should be less susceptible to such differences in the data, but this will be subject to future work. With the current implementation, most of the calculation time is spent for calculating the mutual information of features with selected features. A way to speed this up is by pre-processing and clustering the features so that only a subset of features are subject to mutual information computation for any given feature. The performance of this approach will be evaluated in future work.

## Acknowledgement

We thank Thomas Lengauer, Sarvesh Nikumbh, and Mathieu Flinders for helpful discussions.

